# Single-protein optical holography

**DOI:** 10.1101/2023.08.14.552817

**Authors:** Jan Christoph Thiele, Emanuel Pfitzner, Philipp Kukura

## Abstract

Light scattering by nanoscale objects is a fundamental physical property defined by their scattering cross-section and thus polarisability. Over the past decade, a number of studies have demonstrated single molecule sensitivity, by imaging the interference between coherent scattering from the object of interest with a reference field. This approach has enabled mass measurements of single biomolecules in solution owing to the linear scaling of the image contrast with the molecular polarisability. Nevertheless, all implementations to date based on a common-path interferometer cannot separate and independently tune the reference and scattered light field, prohibiting access to the rich toolbox available to holographic imaging. Here, we demonstrate comparable sensitivity using a non-common path geometry based on a dark-field scattering microscope, similar to a Mach-Zehnder interferometer. We separate the scattering and reference light into four parallel, inherently phase stable detection channels, delivering a five orders of magnitude boost in sensitivity in terms of scattering cross-section over the state-of-the-art, demonstrating detection and mass measurement of single proteins below 100 kDa. Amplitude and phase measurement yields direct information on sample identity and the first experimental determination of the polarisability of single biomolecules.

## Introduction

A core challenge of optical microscopy is to generate detectable image contrast arising from micro-or nanoscopic objects and features. One of the most powerful methods in this regard relies on the phase contrast concept originally introduced by Zernike, which paved the way towards highly sensitive imaging of phase objects. Here, the incident light and that scattered by an object are considered to correspond to the two arms of an interferometer, where the addition of a π/2 phase shift to the scattered field converts a phase shift into an amplitude modulation, which can be imaged.^1,2^ Limitations associated with this initial approach have led to the development of a number of variations of this original concept to enable more quantitative measurements,^3^ and to phase shifting interferometry, where multiple images are recorded to enable combined phase and amplitude imaging.^4-6^ Further optimisation has led to the development of digital holography and quantitative phase imaging, as well as alternative implementations aimed at nanoparticle imaging, tracking, and characterisation.^7-9^ In terms of sensitivity, recent advances have reached metallic nanoparticles as small as 20 nm.^10-13^ Common-path interferometry has been shown to yield holographic information as well,^14^ although it requires recording a sequence of images of the same particle, which limits the sensitivity to particles which are well visible above any static background, such as that of regularly used glass coverslips.^15^

Despite the unique capabilities of these efforts, they have struggled to match the sensitivities that can be achieved with non-holographic, common-path interferometric approaches, which have focused on imaging and quantifying phenomena at interfaces with high sensitivity.^16,17^ The introduction of laser illumination led to a dramatic increase in sensitivity as a result of the combination of higher illumination power densities lowering shot noise and improved spatio-temporal coherence of the illumination. This enabled access to the nanoscale down to 5 nm gold nanoparticles,^18^ and ultimately individual biomolecules.^19,20^ The latter has recently made a substantial impact in characterisation of biomolecular structure, dynamics and interactions through the development of mass photometry.^21-24^ A direct consequence of common-path imaging, however, is that it is difficult to modify the reference field relative to the scattered field leading to a loss of phase information.

Here, we combine sample illumination by total internal reflection with optical quadrature detection,^25^ achieving single-molecule sensitivity reported to date only for common-path approaches. In analogy to a time-domain lock-in amplifier, which mixes sinusoidal waveforms with the modulated signal, we mix four reference fields of the same colour but different phase shifts with the scattered field to retrieve the complex amplitude of the scattered field. Thereby, we measure the complex optical field of sub-diffraction particles, i.e. gold nanoparticles and single proteins. As a result, we can experimentally quantify the polarisability of individual biomolecules, implement holography-specific capabilities such as optimising the focusing in postprocessing,^26^ and demonstrate a step change in the sensitivity achievable with holographic imaging.

## Results

### Proof of concept

Our microscope is based on a Mach-Zehnder interferometer where a scattering object is located in one arm and the reference travels separately along the other arm (Fig. 1a). Placing a quarter wave-plate at 45° in the reference arm generates equal amounts of vertical (v) and horizontal (h)-polarised light with a relative phase shift of π/2. Similarly, a half-wave plate in the scattering arm at 22.5° produces equal amounts of v and h-polarised light with a relative phase shift of π. Recombination by a non-polarising beam splitter (NPBS) generates four phase shifted interferograms: two interferograms in one arm where the horizontal and vertical polarisation contain phase shifts of 0 and π/2, respectively. In the other arm, the two orthogonal polarisations contain interferograms of the phase shifts π and 3π/2. The individual interferograms are then split by two polarising beam splitters (PBS) and imaged onto four individual cameras which are read synchronously.

**Figure 1.**
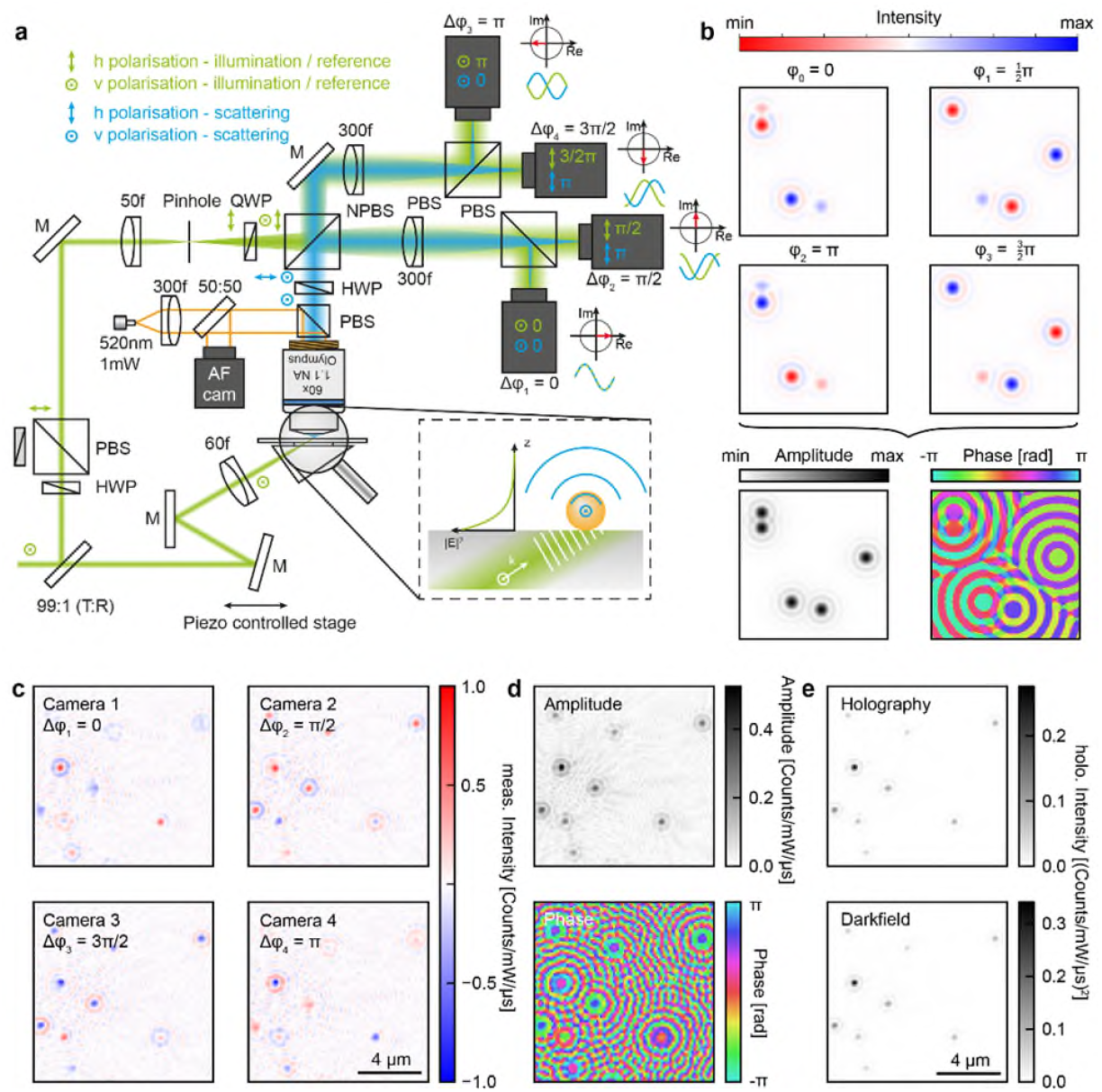
Total internal reflection optical quadrature microscopy. **a** The output from a single emitter laser diode is split by an anti-reflection coated window. One part is focused onto a NBK7 prism to illuminate the sample. The scattered light is collected by a 60x water dipping objective. The other part is mode filtered with a 25 µm pinhole and mixed with the scattered light after introducing different phase shifts to two orthogonal polarisations. Interference of reference and scattered light is deteced for each phase shift by a separate camera. **b** Idealised raw data. Depending on the phase shift, constructive or destructive interference is observed. Combining four images with the correct phase shifts reconstructs the amplitude and phase of the scattered field (lower panel). **c** Representative raw data of 40 nm AuNPs detected by the four cameras after subtraction of any non-interferometric terms. **d** Reconstructed amplitude (top) and phase (bottom) of the electric field. **e** Intensity of 40nm AuNPs derived *via* holography (top) and directly derived by darkfield microscopy.

This approach has several benefits: 1. Total internal reflection illumination enhances the illumination field strength at the interface and suppresses scattering of objects which are further than a few hundred nanometers away from the interface, effectively reducing imaging background. In addition, total internal reflection almost completely extinguishes illumination light reaching the detector, except for that scattered by interface imperfections. 2. The relative amplitude of scattered and reference light can be easily adjusted to match the full well capacity of the imaging cameras at the desired read out speed, as well as to the expected scattering cross sections of the sample. 3. The approach is inherently phase stable, despite a non-common path geometry, because any phase differences identically propagate through all channels, resulting in excellent overall phase stability. 4. Individual blocking of either arm yields the reference intensity, scattering intensity and the interference of both fields separately.

Each camera *j* thus records the interference of the scattered field and the reference field:

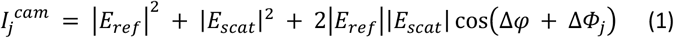

where *ΔΦ*_*j*_ is the additional phase shift introduced by the half and quarter wave plates in front of the NPBS. In other words, each camera records an interferogram of different phase shifts 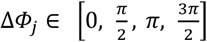 (Fig. 1b), where the scattered light of each particle interferes differently with the reference field in all four cameras. To obtain the desired term, |*E*_*scat*_ | cos(Δ ϕ + Δ*Φ*_*j*_), we subtract any non-interferometric terms from each image (reference field |E_*ref*_ |^2^ and scattering field | E_*scat*_ |^2^) and normalise by |E_*ref*_ |. The reference field and scattered field are recorded directly before measuring the interference of both by blocking either the scattering or the reference path. The resulting images are then combined in a complex fashion according to their phase shifts, by multiplying this corrected output of camera *j* by exp(*i*ΔΦ_*j*_) and computing their average, generating the final complex valued holographic image. This processing is illustrated in supporting figure 1. After complex averaging (real and imaginary part individually), we restore the complex valued scattered field represented by its amplitude and phase (bottom panel of Fig. 1b, and experimental data in Fig. 1c), which allows us to generate the respective amplitude and phase images (Fig. 1d). Reassuringly, the squared intensity derived *via* our holographic approach is identical both in appearance and intensity to the dark-field intensity of the same field of view recorded with identical acquisition parameters (Fig. 1e). These results demonstrate that our approach indeed extracts the correct scattering amplitude, with the only deviation being a constant scaling factor of ~20% likely stemming from imperfect coherence between the reference and scattered fields. This can be accounted for using a calibration if absolute values for the scattered field are desired.

### Demonstration of phase stability

Having verified that we can retrieve the correct scattering intensity, we turn to the precision of our phase measurement. To achieve this, we record movies where we temporally vary the phase between the scattering and reference arm in a linear fashion by a piezo driven mirror in the scattering arm (cf. Fig. 1a). The intensity of individual particles as a function of time reveals clear oscillatory behavior as a function of time (Figs. 2a, b). This is a consequence of the fact that the scattered field of each particle interferes differently with the reference field depending on the camera *j* with a constant phase shift *ΔΦ*_*j*_, and is additionally modulated as a function of time, resulting in a global phase shift *Δϕ* (*t*) between the fields. Following the intensity of a particular particle (marked with a black circle in Fig. 2a) on all four cameras reveals sinusoidal modulations shifted 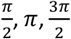 relative to the first camera (blue trace). Despite the fact that the intensity in the individual camera images varies by ±100%, the reconstruction of the complex valued scattered field yields a constant amplitude with a standard deviation of only 0.31%, demonstrating excellent overall phase stability. The phase ramps down linearly as introduced by the movable mirror. The same behavior can be observed for all other particles in the region of interest (grey traces in Figs. 2c, d).

**Figure 2.**
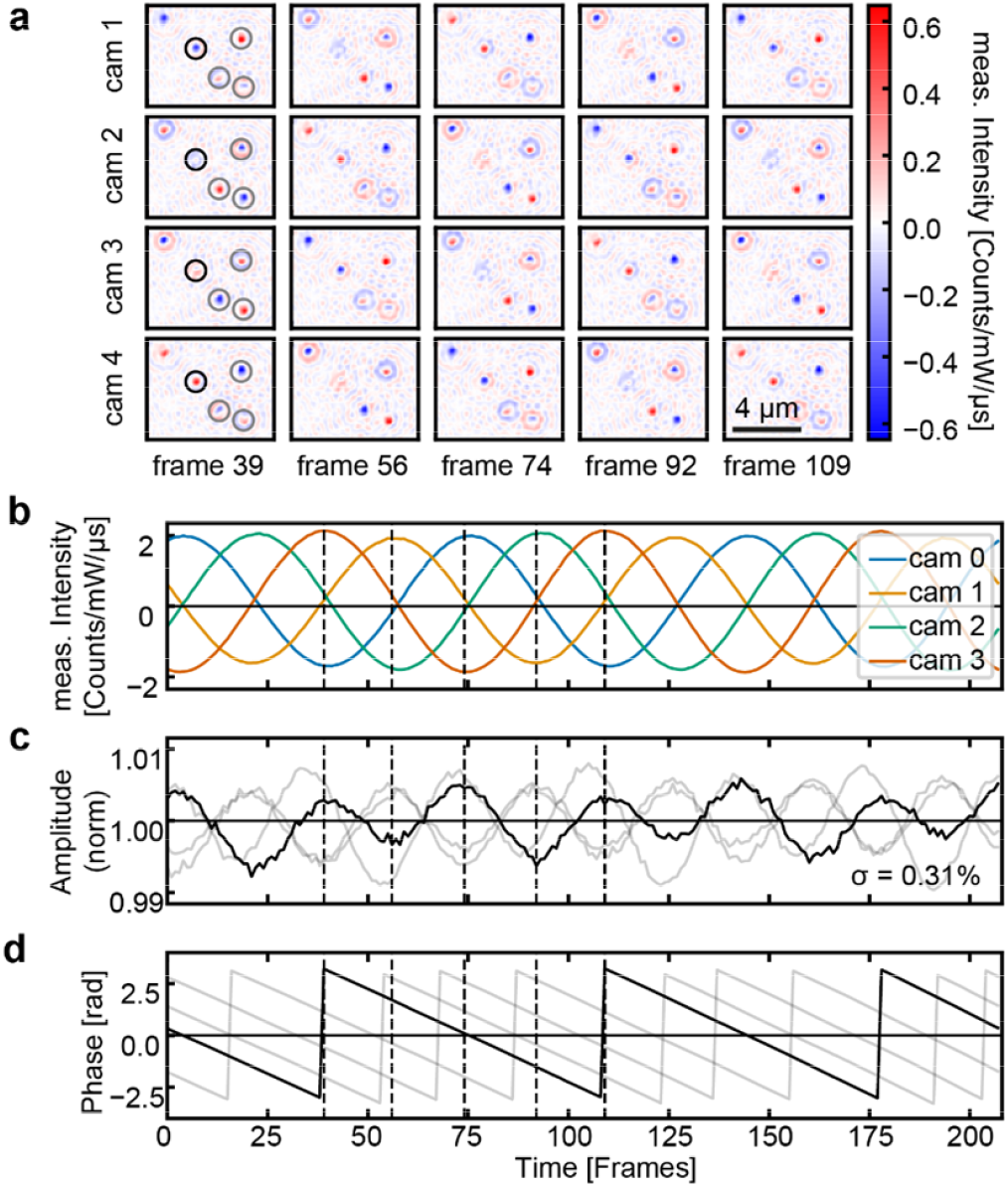
Holography of 40 nm AuNPs and resulting phase stability. **a** Images of four cameras (phase shifts 0, π/2, π and 3π/2) with different phase shifts introduced by a movable mirror in the illumination arm. **b** Temporal evolution of the detected interference of a single AuNP (black circle in a). Each trace corresponds to a different camera. Reconstructed amplitude (**c**) and phase (**d**). The black trace corresponds to the highlighted AuNP, the gray colored ones to the other particles in the field of view.

### Optical holography of individual proteins

Given these encouraging results, we queried to which degree this approach may be suited to detect, image and quantify very weak scatterers, such as individual proteins. A cleaned microscope glass coverslip produces a speckle pattern (Fig. 3a) reminiscent of that observed in iSCAT^27^ and high performance dark field microscopy.^28^ Again, the darkfield and holographically reconstructed images show an excellent quantitative agreement. Introducing ~40 nM of a soluble 90 kDa protein (DynΔPRD) results in no discernible changes to the amplitude images as a function of time (Fig. 3b). Computing the differences of the mean of two moving subsequent windows of length 10 frames (6.25 ms/frame), however, reveals clear signatures of individual proteins binding to the surface in both amplitude and phase (Fig. 3c). We then quantified the contrast of each landed protein by fitting a complex valued point spread function (PSF) model derived by the average PSF of all landing events (Fig. 3d). The resulting histogram exhibits several sub-distributions as expected for the oligomeric nature of dynamin (DynΔPRD), corresponding to the monomer, dimer, tetramer and hexamer. The fitted masses of the first four distributions correlate well with the expected molecular weights of 90, 180, 360, and 540 kDa (Fig. 3e). The linear relationship between contrast and molecular weight allows us to deduce the mass of any unknown protein based on its image contrast.

**Figure 3.**
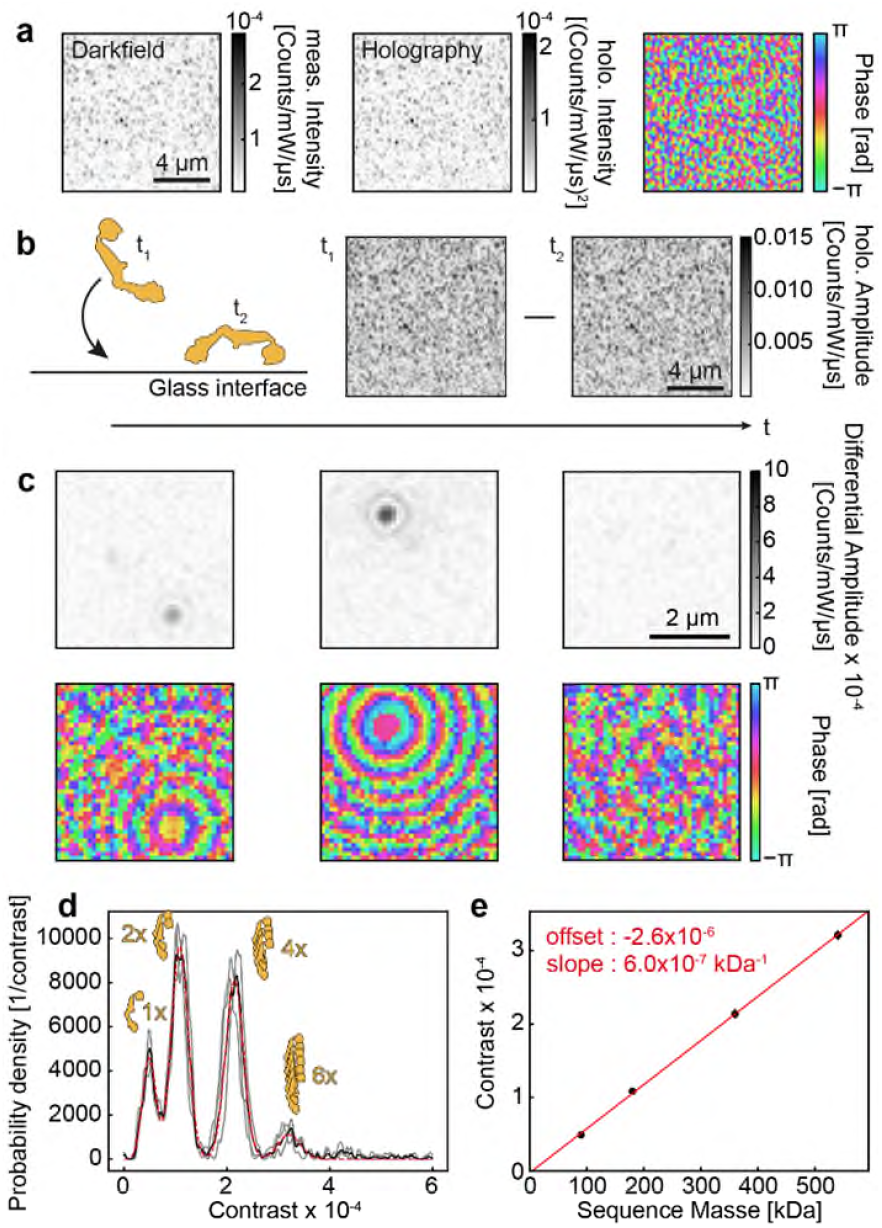
Single-protein holography. **a** Darkfield and holography reconstructed intensity and phase maps of a microscope coverslip exhibiting typical residual glass roughness. **b** The amplitude of the glass does not change visibly before and after a single protein landed due to its low scattering cross-section. **c** Moving mean differences of subsequent image stacks reveal individual proteins landing in the amplitude and phase representation. **d** Amplitude histograms (grey) of the landing events from individual movies acquired over 120 seconds. The red line represents a fit of four gaussians to the averaged data shown in black. **e** Fitted contrast vs sequence mass of the different oligomers. The red line indicates a linear fit to the data points. Error bars are in the order of the data point size.

In principle, we could also derive the scattering intensity by collecting the darkfield scattering as shown in figure 1e. Supporting figure 2 shows a histogram of protein landing events at the same glass spot generated via holography and synthetic darkfield. The synthetic darkfield histograms, however, exhibit a much lower mass resolution. Comparing individual PSFs illustrates why the resolution decreases: only in the holographic approach are the PSFs well-defined whereas each individual PSF looks different for darkfield imaging. This is due to the fact, that in darkfield operation the scattered field interferes with a the complicated wavefront of the light scattered by the glass interface, which prevents an accurate estimate of the contrast.

Until now, we only investigated the amplitude of the landing events. However, we simultaneously derive the phase of each event. Due to the oblique illumination associated with total internal reflection, we imprint a linear phase gradient onto the illumination field (Fig. 4a)

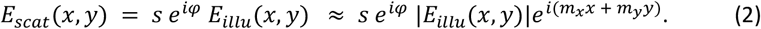

**Figure 4.**
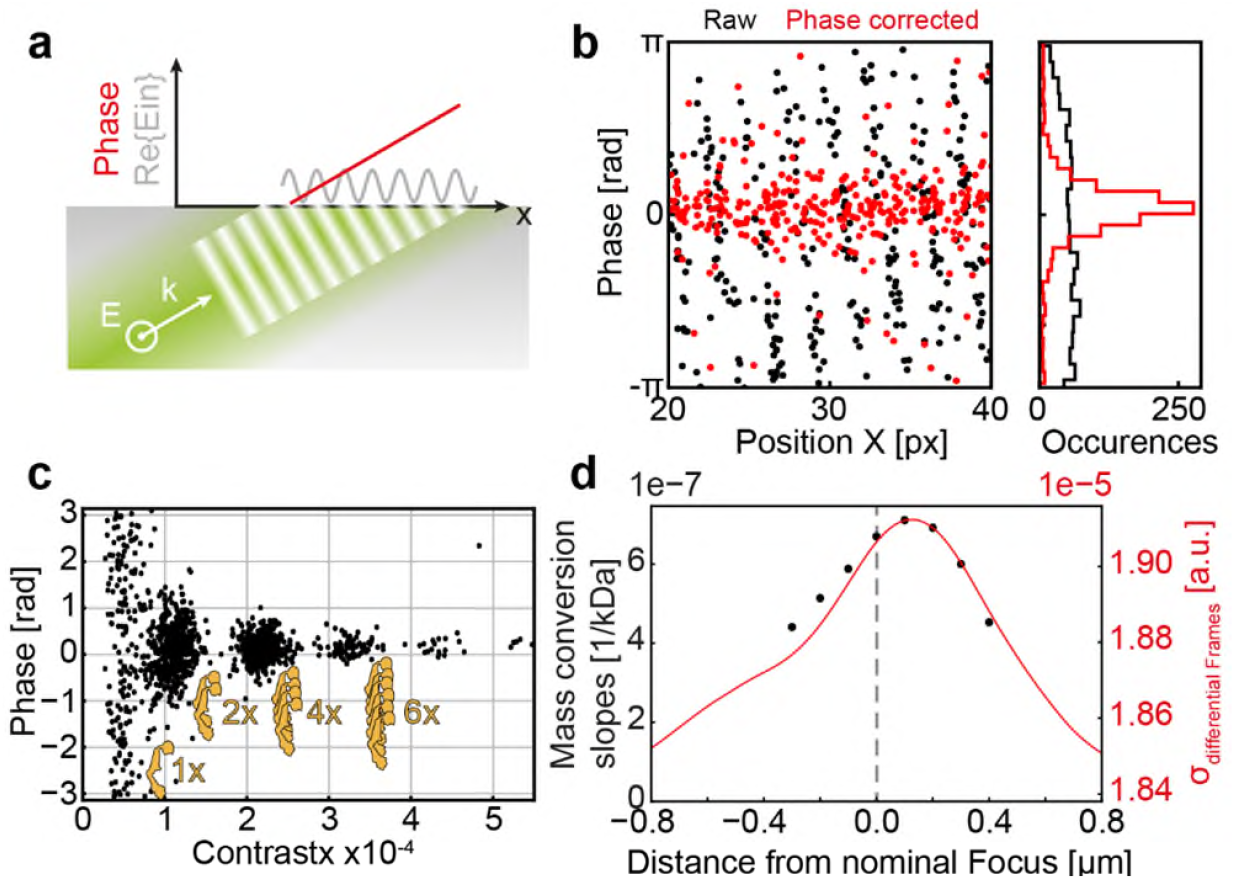
Phase-sensitive microscopy of single-proteins. **a** Oblique angle of illumination imprints a phase gradient onto the scattered light as a function of landing position. **b** Phase values of landing events vs landing position (black data points). Subtraction of a linear phase gradient removes the spatial dependence of the phase (red data points). **c** Phase of each landing event vs its contrast. **d** Contrast to mass conversion slopes (black data points) vs post-processing focus. The underlying contrast histograms are provided in supporting Fig 4a. The black line shows the standard deviation of the differential frames for different post-processing focus positions.

Where s and *ϕ* are the intrinsic scattering amplitude and phase of the protein, *E*_*illu*_(*x, y*) the illumination field, and *m*_*x*_ and *m*_*y*_ the phase gradient along *x* and *y* due to the oblique angle of incidence. Inspection of the phase difference of all landing events as a function of their position along the illumination direction reveals a linear correlation (supporting figure 3). We make use of this linear correlation to find a slope *m*_*x*_ along the illumination direction and *m*_*y*_ perpendicular to it which removes the linear phase gradient. We then subtract a linear phase of each landing events phase *ϕ*_*i*_ such that

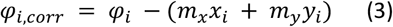

where *x*_*i*_ and *y*_*i*_ denote the landing position of the landing event *i*. Applying this correction to all landing events globally (Fig. 4b, black data points) shifts the phases of almost all events onto a straight line (Fig. 4b, red data points). The corrected phase of all individual oligomeric species falls onto a line as expected for a non-absorbing dielectric nanoparticle (Fig. 4c).^11^ The corrected phase becomes less well defined the lower the contrast of the particle. Supporting figure 6 suggests, that the localisation precision dramatically decreases with lower contrast. This affects the phase correction which utilises the position of the landing event. Note that we are not able to derive the absolute phase of the landing particles as we lack knowledge about the phase relation between the illumination field and the reference field. The lack of any correlation of the phase with the event contrast further underpins the separation of amplitude and phase of our approach. Knowledge of the complex scattered field enabes us to change the focus during analysis of the data (cf. SI Fig. 4, see Supporting Information for more details).^29^ The standard deviation derived form the differential amplitude data correlates excellently with the contrast to mass conversion (Fig. 4d). We make use of this to automatically optimise the focussing conditions for all our measurements allowing us to maximise the imaging contrast in post processing.

### Measurement of the polarisability of a single protein

Our holographic approach quantifies the scattered field independently of any unknown phase and optimises focussing during post-processing, both for AuNPs and dielectric proteins. In the following, we use these capabilities to derive an experimental estimate for the polarisability of a protein attached to a glass coverslip per molecular weight which we will call specific polarisability. As a calibration, we first measure the contrast of static AuNPs for different sizes (20 nm, 40 nm, and 60 nm). The contrast is normalised to the exposure time and laser power which we both reduced for the larger AuNPs (40 nm and 60 nm). We did not use smaller AuNPs to ensure that they are clearly visible above the coverglass background. Plotting the third root of the contrast for the holographic data (Fig. 5a) and the sixth root of the contrast for the darkfield contrast (Fig. 5b) linearises the plotted quantity with respect to the particle volume as long as Rayleigh scattering dominates. In both cases – holographic and darkfield – the mean contrast increases linearly with c^1/3^ and c^1/6^, respectively, which again suggests that the retrieved scattering amplitude is in agreement with theoretical predictions. The values gathered *via* darkfield microscopy are 20% (in intensity) and 3% (in intensity^⅙^) larger than the values retrieved by holography in agreement with imperfect coherence as discussed in Fig. 1e.

**Figure 5.**
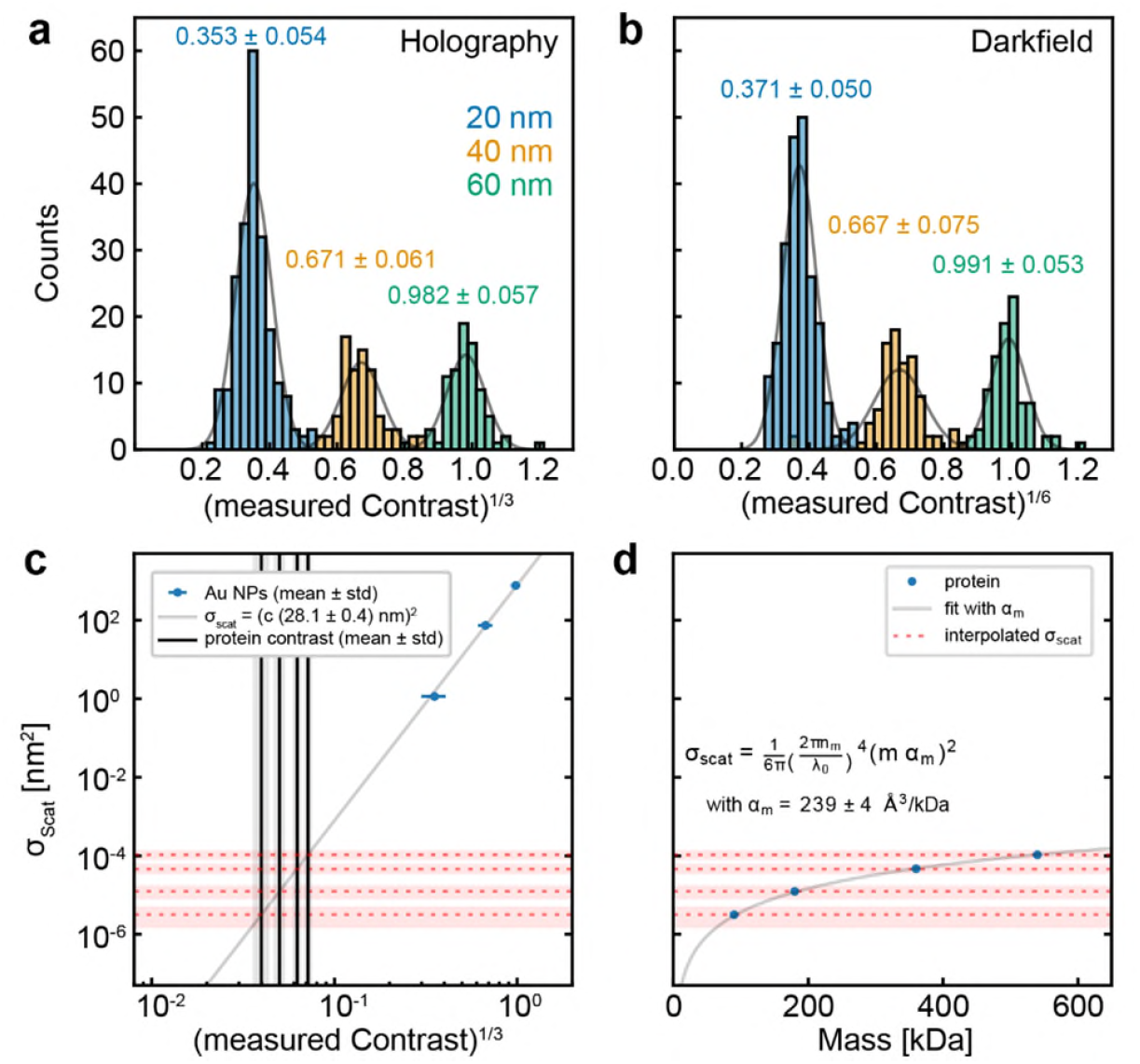
Quantification of the polarisability of a single protein. **a** Histogram of the third root of the holographically derived particle contrast and **b** sixth root of particle contrast derived from darkfield images for 20 nm, 40 nm, and 60 nm AuNPs. **c** Calibration of the scattering cross-section using AuNPs. The blue data points are the experimentally derived contrasts vs their scattering cross-section calculated using Mie theory. The gray line is a one parameter linear fit to the 6^th^ root of the scattering cross-section. The black lines indicate the measured contrast of our oligomeric protein sample. **d** Based on the calibration in **(c)**, we can plot the scattering cross-section of protein (red dashed lines) vs their theoretical molecular weight (blue data points). The solid line represents a fit of the eqn. 4 to the data.

For each particle size, we calculate a scattering cross section (Fig. 5c) by assuming the AuNPs to be homogeneous and embedded in a homogeneous medium of refractive index n = 1.337. As previously shown,^30^ the scattering cross-section of the herein used AuNPs ranging from 20 to 60 nm can be well estimated by Mie scattering. This allows us to perform a one-parameter linear regression on the derived scattering cross-section against our measured contrast. We are now in the position to estimate the scattering cross-section of the measured protein assuming that the collection efficiency and surface related enhancement effects are identical for AuNPs and proteins (Fig. 5d). We assume the scattering of the protein to be well described by Rayleigh scattering of a dielectric particle:^31^

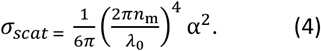

Fitting the scattering cross-section of the protein to equation 4 yields a specific excess polarisability *α* per molecular weight of 239 ± 4 Å^3^/kDa. The statistical error is based on error propagation from the standard deviations of the fit parameters. Assuming a specific volume of 0.7446 mL/g,^22,32^ we can estimate the refractive index of protein to *n*=1.47. Both these values for the specific polarisability and the refractive index are well within the range of values reported previously in the literature,^33-35^ although somewhat smaller than that recently calculated using an atomic polarisability model.^36^ We remark that this polarisability measurement by comparison with AuNPs is distinct to similar efforts made with a common path geometry, such as in iSCAT or mass photometry, because they only yield the intensity, convolving amplitude and phase contributions, the latter of which is generally not well-defined.

## Discussion

Using an inherently phase stable approach, we have demonstrated holographic imaging with a sensitivity reaching single biomolecules in solution. While in its current implementation, the approach was unable to match the sensitivity and measurement precision achievable with mass photometry, there are clear routes to potential future improvements. First and foremost, our current experimental arrangement was limited to a power density of 50 kW/cm^2^, an order of magnitude lower than that used in mass photometry and ensuing loss in shot noise limited sensitivity. In addition, we did not yet optimise the electric field enhancement due to total internal reflection as a function of incidence angle, resulting in a relatively low 10% enhancement compared to the achievable factor of three.

We thus expect a potential order of magnitude improvement in sensitivity and thereby mass resolution, which would surpass that of all current single molecule detection approaches. These advantages are likely to be compounded by improved background rejection stemming from total internal reflection known from fluorescence microscopy. We further expect improvements from an optimised illumination source in terms of spatio-temporal coherence and frequency stability over that achievable with a low cost laser diode used here. Finally, illumination with a precisely pre-tilted wavefront via reflection off a suitable grating would compensate for the positional dependence of the scattering phase and reduce the uncertainty caused by limited localisation precision in the current implementation. Additionally, a bleed-through channel via the scattering arm would enable measurements of the scattering phase in absolute terms.

Taken together, we have presented an implementation of optical quadrature microscopy in combination with total internal reflection illumination capable of holographic imaging of single biomolecules. We verified our approach by showcasing the ability to separate the amplitude and phase of the scattered field and confirmed its quantification by direct comparison with darkfield microscopy. The resulting sensitivity leads to single protein imaging, which can be converted into a measurement of the the polarisability of a single protein based on the scattering of a well-characterised reference object. These results break new ground for optical holography, providing access to the exciting sub 20 nm length scale for phase and amplitude imaging with applications in the biological, physical and material sciences.

## Methods

### Setup

The optical setup used in this study is similar to previously reported optical quadrature microscopes.^25,37^ A single-emitter laser diode (450 nm, 500mW, NDB4816, Nichia) is mounted in a temperature-controlled mount (Thorlabs GmbH, 400 mA, 17 °C) and collimated using an aspheric lens (f = 3.30 mm, N414TM-A, Thorlabs LTD.). The polarisation is aligned vertically using the light reflected off a PBS (PBS251, Thorlabs LTD.) and guided to an anti-reflection coated NBK7 window (WW11050-A, Thorlabs LTD.) which picks up ~0.5 % of the incident power and reflects it into the reference arm. >99% of the light propagates towards the prism (12.7mm Littrow Prism (90-60-30), Edmund optics) and is focused into the focal plane of the objective using an achromatic lens (f = 60 mm, AC254-60-A, Thorlabs LTD.). Standard glass coverslips (24×75 mm, thickness 1.5, Carl Roth) are optically coupled to the prism by immersion oil (n=1.52) to enable fast exchange of samples without the need for tidiously cleaning the prism. Scattered light from the coverslip surface is collected using an objective (60×, 1.1NA, water dipping, Olympus) and is imaged onto four different cameras by an achromatic tube lens (f = 300 mm, AC254-300-A, Thorlabs LTD.). It is important to note that the tube lens is placed one focal length apart from the back focal plane of the objective to create a flat wavefront in the image plane. The light reflected towards the reference arm is focused by an achromatic lens (f = 50 mm, AC254-50-A, Thorlabs) onto a 25 µm pinhole (P25CB, Thorlabs LTD.) to filter the mode of the laser. The diverging light is then collimated by the tube lens.

The central elements of this optical scheme are the two waveplates in front of the non-polarising beam-splitter (NPBS): the two orthogonal polarisations of the reference arm are phase shifted by the quarter waveplate resulting in a phase shift of 0 and π/2 for the horizontal and vertical polarisation, accordingly. External reflection off the NPBS introduces an additional phase-shift of π whereas the transmitted polarisations are not affected. Light coming from the scattering arm propagates through the half-wave plate which introduces 0 and π phase shift to the two orthogonal polarisations. Mixing the reference and scattering light at the NPBS and separating them via a PBS allows to create an image with the relative phase shifts of 0, π/2, π, and 3π/2. The four cameras (Grasshopper GS3-U3-23S6M-C, Teledyne FLIR) detect the four phase shifted images at 800 frames per second in a synchronised fashion. Five consecutive frames are averaged and the data of all cameras is aligned to the first camera (affine transformation: scale, shift, rotation) and 2×2 pixels are binned resulting in an effective pixel size of 117.2 nm.

To find and maintain focus throughout all measurements, a collimated laser beam (0.5 mW, 520 nm, Thorlabs LTD.) propagates through the back aperture of the objective. The light is focused onto the coverslip-buffer interface. The reflected light is collimated and imaged by a camera through a 50/50 non-polarising beamsplitter. The radius of the resulting circle is maintained by a PID controller which controls the axial distance between objective and coverslip. All data acquisition and control is handled by a custom-built LabView program.

### AuNP samples

All gold nano-particles were purchased from NanoPartz Inc. (A11-20-CIT-DIH-1-10, A11-40-CIT-DIH-1-10, and A11-60-CIT-DIH-1-10). First, clean coverslips were rendered hydrophilic by placing them in an O_2_ plasma for 30 s (30% power, 0.5 mBar O_2_, Zepto-BRS 200, Diener electronic GmbH). For all sizes of gold Nanoparticles, 10 µl were spin-coated onto the hydrophilic coverslips for 18 s at 500 rpm and 18 s and 60 s at 1400 rpm. A PDMS gasket (CultureWell CW-8R-1.0, Grace Bio-Labs) was placed at the center of the coverslip and 50 µl milliQ water added to achieve immersion with the objective lens.

### Protein landing

DynΔPRD^23^ was diluted 25-50 fold from its stock concentration to 80 – 40 nM before replacing the droplet for immersion with the diluted solution. Landing events were recorded for 2 minutes after replacing the immersion solution.

### Data analysis

All analysis was performed in custom python scripts. Raw data was first aligned by finding an affine transform which minimises the least squares of the differences of the aligned and target image. We used only the scattered intensity for aligning the four channels since it showed higher contrast than the interferometric measurement and resulted in a more reliable alignment. The data was then pixel binned (2×2 binning). To extract the complex scattered field of static images, first the reference image and the darkfield (both recorded shortly before acquiring the holographic data) were subtracted. To extract the scattered field only, the resulting image was divided by the square root of the reference image. The resulting image now only contains 2|*E*_*scat*_ | cos(Φ_*j*_) projected along the positive and negative real and imaginary axis of a complex plane. By averaging the four images rotated all onto the positive real axis, recovers the complex field E_*scat*_.

For static measurements, particles were detected on the absolute of the reconstructed scattering field |*E*_*scat*_ | by calculating a radial symmetry map (α=1.0, radius=3 px)^38^ and finding local maxima with radial symmetry above the 99 % quantile of the map.

For fitting the scattered field of particles, the following complex valued point spread function model was used:

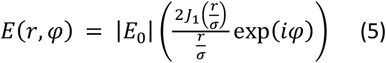

Where *J*_1_ denotes the Bessel function of first kind, *σ* the width, *r* the distance from the center, |*E*_0_| the amplitude, *ϕ* the phase of the landing event. The squared residuals (*SR*) of the sum of the real and imaginary parts of the point spread functions were used as an objective function for minimisation by optimising *x, y*, |*E*_0_|, and *ϕ*:

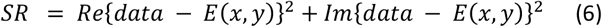

For the data presented in Figure 2c, 2d, and 5a, measurements of immobilised nano particles were analysed during which the length of the sample path was varied with a piezo shifted mirror to introduce a phase shift. A segment of the measurement where the phase shift linearly increased by 6π was selected, particles were detected on a sum of |*E*_*scat*_ | over all phase shifts. A 23x23px thumbnail was fitted with the complex PSF for each phase shift independently. For Figure 5a, the cubic roots of the amplitudes were averaged over all phase shifts and particles with a relative standard deviation of the cubic root of the amplitude above 10% or a standard deviation of the position above 0.2 px were rejected.

The darkfield measurements of immobilised nano particles were analysed in a similar way. For the particle fitting, the scattered intensity |*E*_*scat*_ |^2^ was fitted with an intensity PSF (square of the complex PSF in eqn. 5) in all four channels independently. For Figure 5b, the sixth roots of the amplitudes were averaged over all channels and particles with a relative standard deviation of the sixth root of the amplitude above 10% or a standard deviation of the position above 0.2 px were rejected.

For dynamic movies (i.e. landing events of proteins), only the reference image was subtracted. The resulting image was divided by the square root of the reference and the complex movie was calculated. Temporal phase changes were corrected by applying the phase shift which minimises the difference to the first frame. The corrected complex movie was then further analysed by a sliding difference of 10 frames window length to remove static features like the glass roughness and other proteins which landed before the analysed window. As shown in supporting figure 7, the phase correction reduces the noise in the differential images.

For detecting the position and time of the landing events, a normalised cross-correlation with a complex PSF (eqn. 5) was applied to the movie followed by a maximum filter. The resulting mask was then multiplied by the previous normalised cross-correlation and thresholded at 0.15. This essentially gives a first approximation of the location and time of all landing events. We then convolved the mask with a circle of radius 2 along the spatial axes and a pulse function along the temporal axis of length 10 frames to blur the found particle locations. We then searched for the local maximum in this mask which we define as landing position and time. A 23px x 23px thumbnail at the detected landing event position was fitted to the complex PSF as described above to find exact spatial positions. These thumbnails were used to generate a an experimental PSF by interpolating the average of all aligned thumbnails. In a second step, the fits were refined by fitting with an experimental PSF. The events contributing to the histogram were filtered according to positional precision (fit position closer than one pixel to initial guess), fit position (omit events too close to the border and trim events where the illumination intensity is too uneven) and fit residuals (cf. supporting figure 5).

### Calculation of the protein polarisability

The scattering cross-section of the gold nanoparticles was calculated with the Mie formalism^39^ for a wavelength of 450 nm and a refractive index of n_Au_=1.528+1.911i for gold^40^ and 1.337 for the embedding medium (H_2_O).^41^ The linear scaling factor between the cubic root of the interferometric contrast *c* of the single particles and the sixth root of the scattering cross-section *σ*_*scat*_ was determined with an ordinary least squares^42^ *σ*_*scat*_ = (c * (28.1 ± 0.4)nm)^2^. By using the cubic root of the interferometric contrast, a normal distribution in particle diameter is assumed instead of a normal distribution in volume. With this relation, the scattering cross-section of the different protein oligomers shown in Fig 5d is estimated. From the Gaussian fits to the contrast distributions of the oligomers and their known masses *m*, the relationship c = *m*(6.82 ± 0.04)10^−7^/kDa was determined with a least square regression using the invers of the fitted peak variance as respective weights. Combining these two relationships with the formula for Rayleigh scattering of a dielectric particle in eq. (4) yields a specific polarisability 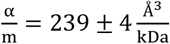.

## Acknowledgement

We thank the Micron Bioimaging Facility of the University of Oxford for a loan of water immersion objective and Dr. Jan Becker for fruitful discussions. The protein was supplied by Dr. Manish Kushwar.

Emanuel Pfitzner was supported by the Deutsche Forschungsgemeinschaft (DFG, German Research Foundation) under project 455633413.

The work was supported by European Research Council (ERC) Consolidator Grant PHOTOMASS 819593 and Engineering and Physical Sciences Research Council (EPSRC) Leadership Fellowship EP/T03419X/1 (P.K./J.C.T). For the purpose of Open Access, the author has applied a CC BY public copyright licence to any Author Accepted Manuscript (AAM) version arising from this submission.

