## Supplementary information for "Single-protein optical holography"

### Supplementary information:
### Single-protein optical holography

Jan Christoph Thiele*, Emanuel Pfitzner*, and Philipp Kukura

*The Kavli Institute for Nanoscience Discovery, University of Oxford, Dorothy Crowfoot Hodgkin Building, South Parks Rd, Oxford OX1 3QU, U.K.*
*Physical and Theoretical Chemistry Laboratory, Department of Chemistry, University of Oxford, South Parks Road, Oxford OX1 3QZ, U.K.*
*These authors contributed equally.

---

#### Refocussing

To refocus the complex image, a frequency-dependent phase shift is applied in Fourier-space: The phase shift is defined as

$$\Delta\varphi\left(k_x, k_y\right) = \Delta z\, k_z\left(k_x, k_y\right) = \Delta z \sqrt{k^2 - k_x^2 - k_y^2}$$

with $k$ the wavevector in the medium (water) and $\Delta z$ the focus offset. For frequencies above the support ($k^2 < k_x^2 + k_y^2$), the phase shift is set to 0. To apply the phase shift, the image is Fourier transformed, multiplied by $e^{i\Delta\phi}$, and back transformed to real space.

#### Angle of incidence

We used the slope of our phase correction to determine the angle of incidence. The phase correction of about 2.4 rad / px can be easily transferred into 48.83 nm / rad utilising the total magnification of 100x given by the focal lengths of the used lenses. This translates to 306.8 nm / wave at the interface. Using trigonometry we can estimate the angle of incidence to 74.8 deg assuming the vacuum wavelength of 450 nm and a refractive index of n = 1.52 for the coverslip.

### Supporting Figures

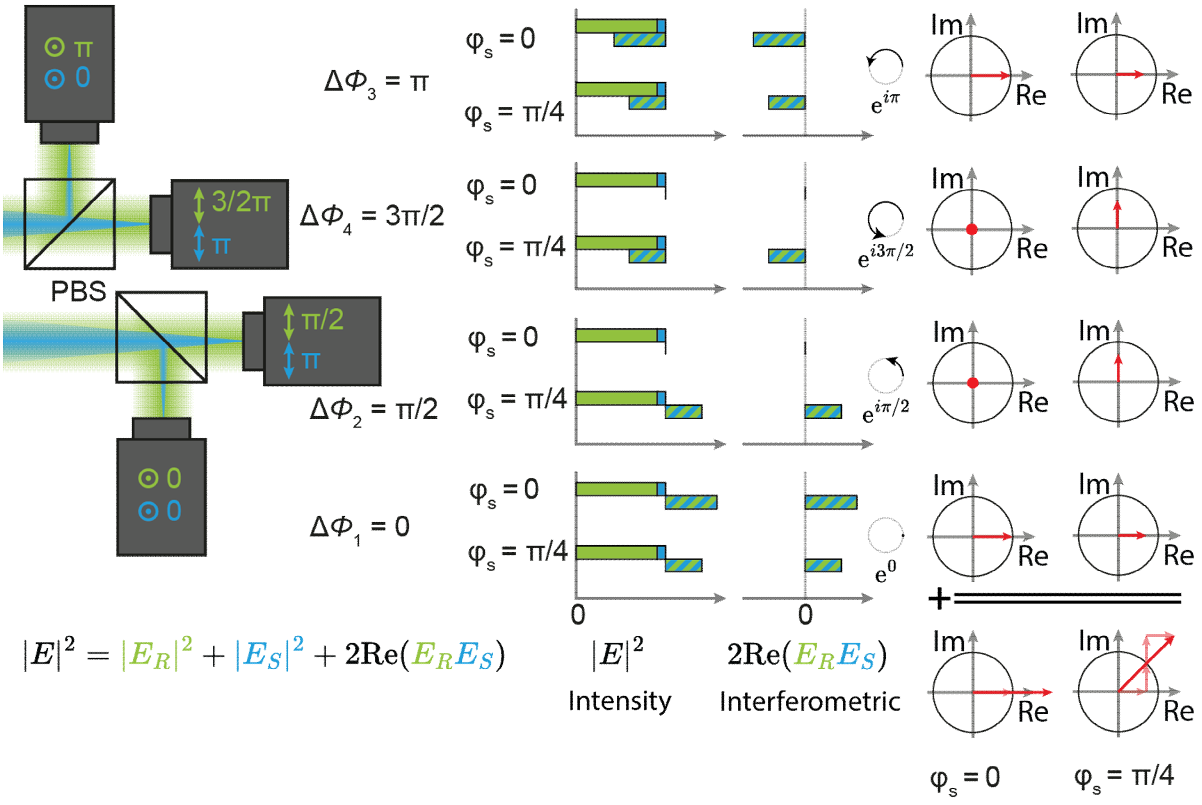


**Supporting Figure 1 – Reconstruction of holographic images.** All cameras receive light with a different relative phase shift between the reference field $E_R$ and the scattering field $E_S$. The interferomtric component is calculated for each camera by subtracting the prerecorded reference intensity $|E_R|^2$ and scattering intensity $|E_S|^2$. The unnormalised complex scattering field is otained by summing all interferomtric components after correcting for their respective phase shifts.

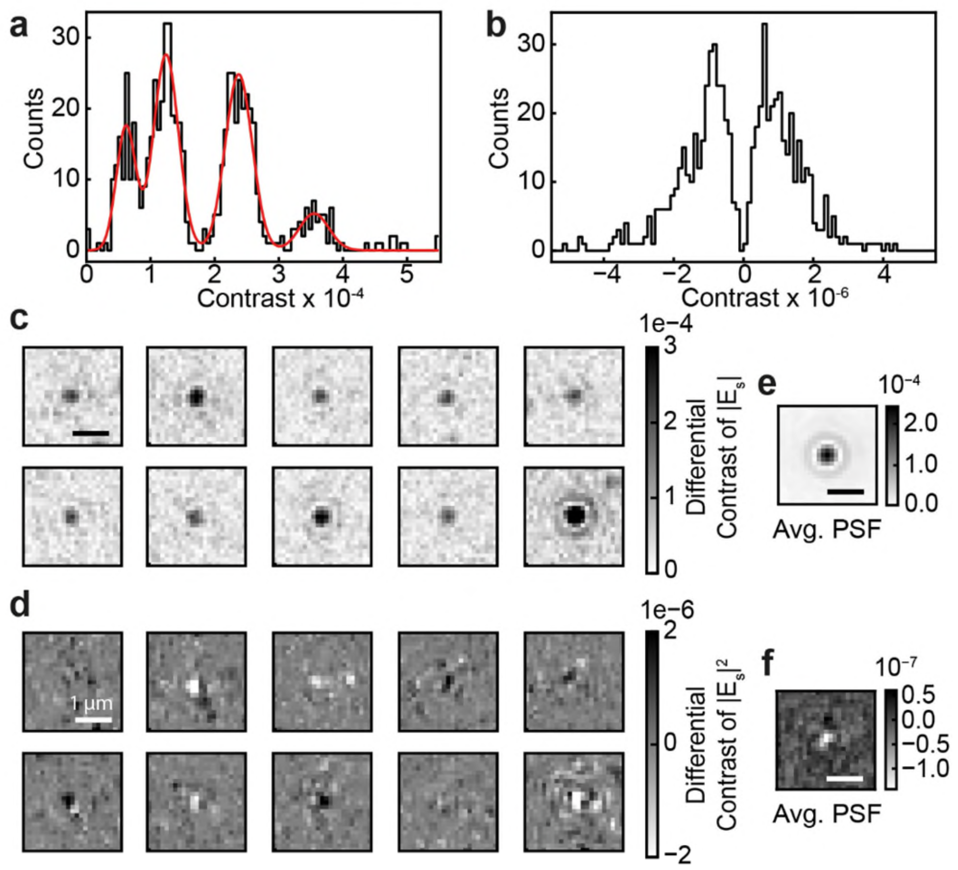


**Supporting Figure 2 – Comparision of holographic and synthetic darkfield contrast histogram.** **a** Histogram of the landing event contrast determined by fitting the scattered field $E_{scat}$ with a complex, holographic PSF of one movie. **b** Histogram of the contrast from the same landing events fitted in a synthetic darkfield movie. The darkfield movie was generated by calculating the square of the absolute of the scattering field $|E_{scat}|^2$ where $E_{scat}$ was derived by our holographic approach. **c** Example PSFs of landing events in the holographic movie and **e** the average PSF of all fitted events. **d** Example PSFs of the same landing events in the synthetic darkfield movie and **f** the average PSF.

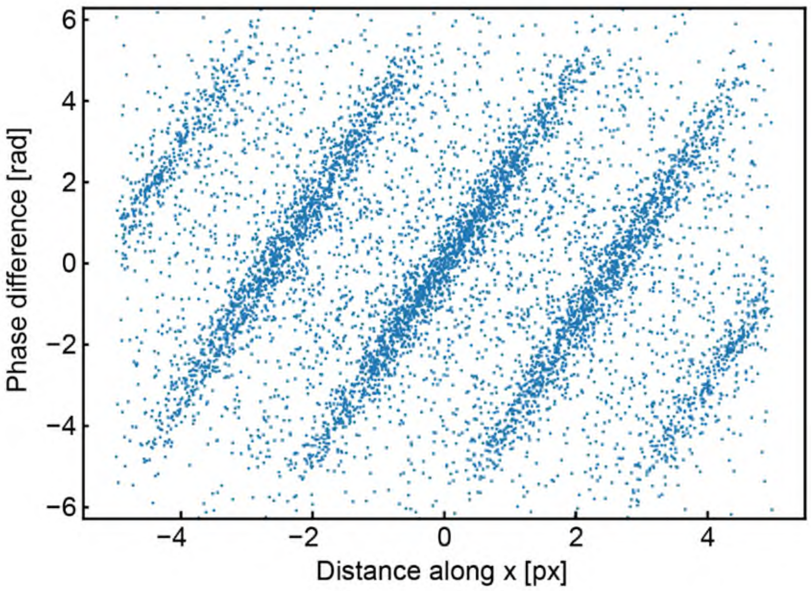


**Supporting Figure 3 – Positional dependence of the phase.** Difference of the detected phase $\Delta\varphi$ of individual landing particles vs their distance in landing position along the illumination direction. The linear correlation of $\Delta\varphi$ and distance relates to the oblique angle of illumination.

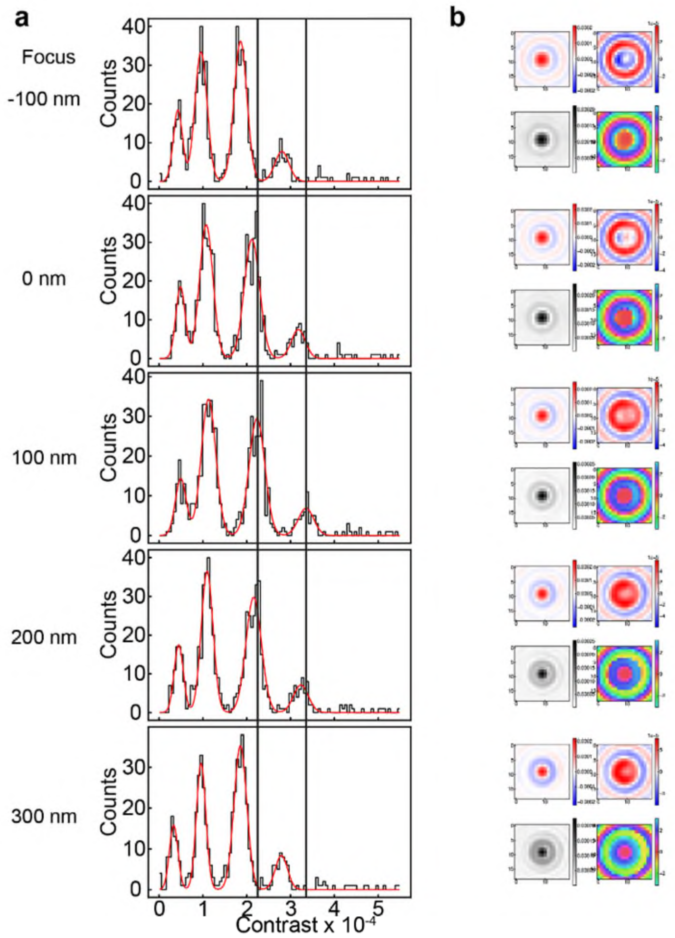


**Supporting Figure 4 – Single-protein contrast vs focussing during post-processing**. **a** Histograms fitted with experimental PFSs for different focus positions. The red line corresponds to a sum of Gaussian distributions fitted to the histograms. **b** Real-part, imaginary-part, amplitude and phase of average point spread function model used to fit the data.

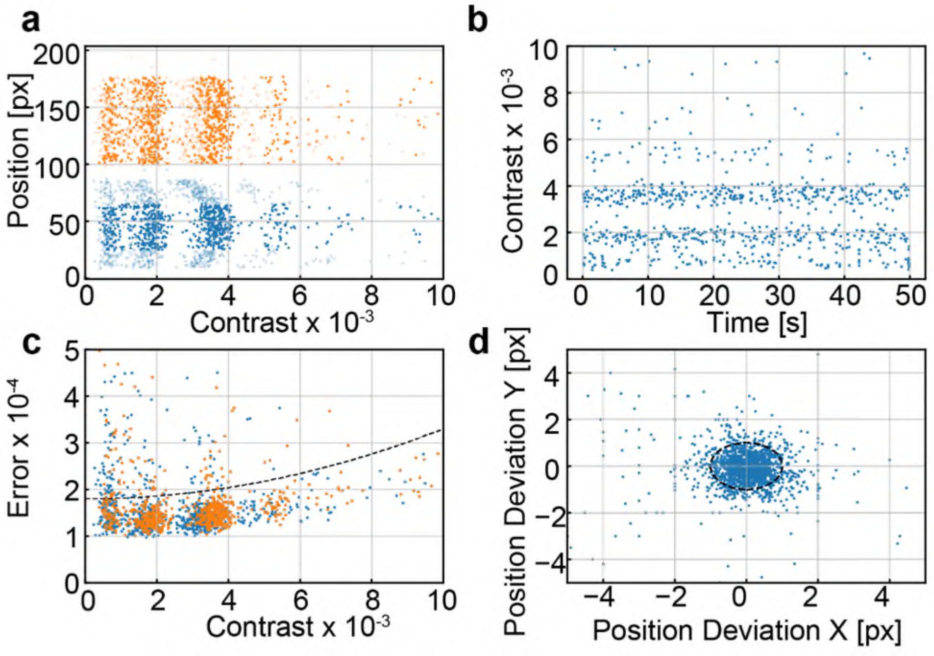


**Supporting Figure 5 – Selection criteria for protein landing events. a** Contrast vs x position (orange) and y position (blue) of each landing event. Only events in the center along y axis (solid blue dots) are selected. **b** Temporal evolution of the selected events confirm axial stability during an acquisition. **c** Residuals of all fits. Events above a quadratic threshold (dashed black line) are neglected similar to Foley, et al. [23]. **d** Deviation of fitted position from initial guess. Events which deviate more than 1 pixels from the initial guess (black dashed line) are discarded.

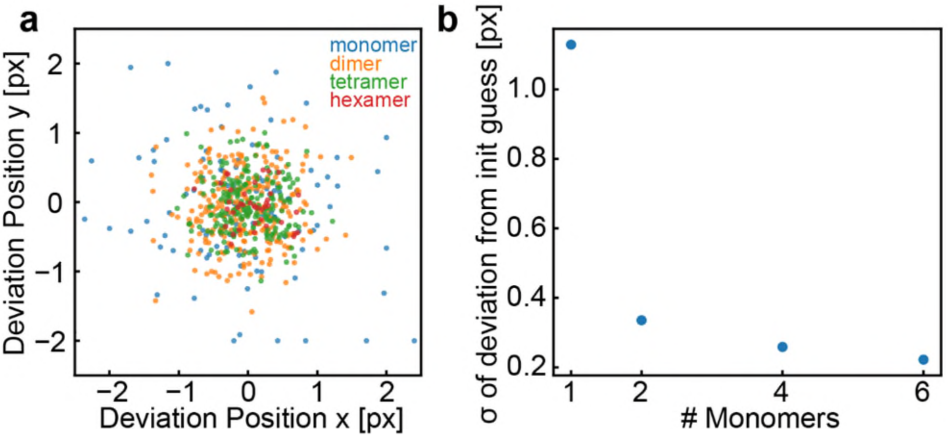


**Supporting Figure 6 – Localisation precision. a** Deviation of fitted position from initial guess. **b** width of distributions in **a**.

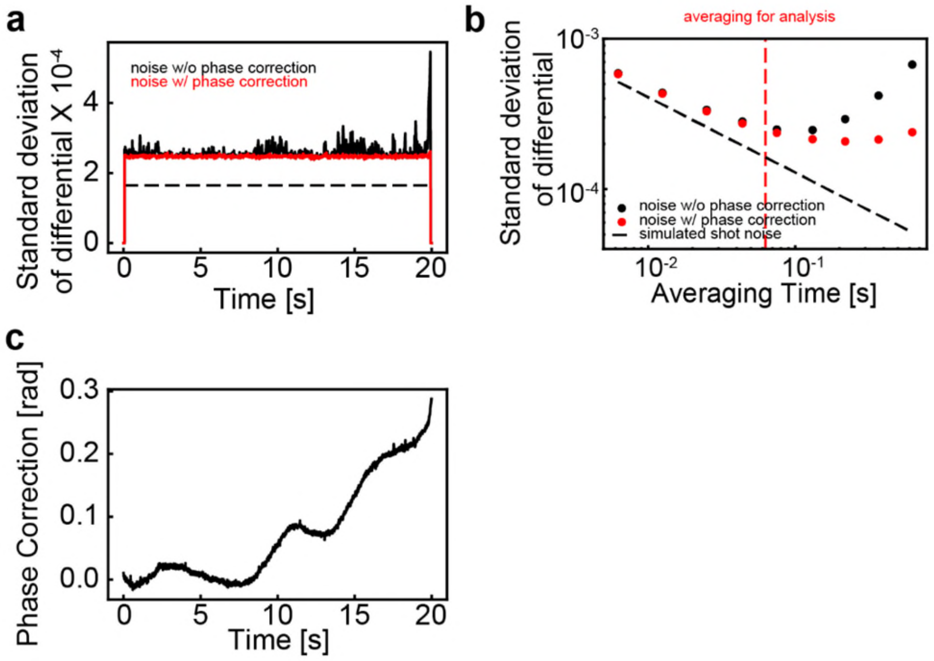


**Supporting Figure 7 – Signal-to-noise. a** Noise vs time of the differential of the complex valued scattered field for a movie containing no protein (buffer only). The red line indicates the noise trace after applying a phase correction on a frame-by-frame basis. **b** Standard deviation in the differential frames vs averaging time without phase correction (black data points) and with phase correction (red data points). The red dashed line indicates the averaging time used for the analysis of protein landing events. The black dashed line indicates the shot noise of a simulated movie with the same amount of detected photons per unit time compared to the experimental data. **c** Temporal phase evolution to correct data in **a** and **b**.
